# PgtE protease enables virulent *Salmonella* to evade C3-mediated serum and neutrophil killing

**DOI:** 10.1101/2024.11.05.622138

**Authors:** Michael H. Lee, Araceli Perez-Lopez, Leigh A. Knodler, Grace Nguyen, Gregory T. Walker, Judith Behnsen, Steven Silva, Jean Celli, Melissa A. Tamin, Michael H. Liang, Karine Melchior, Felix A. Argueta, Sean-Paul Nuccio, Manuela Raffatellu

**Affiliations:** Division of Host-Microbe Systems and Therapeutics, Department of Pediatrics, University of California San Diego, La Jolla, CA 92093, USA; Biomedicine Research Unit, Facultad de Estudios Superiores Iztacala, Universidad Nacional Autónoma de México. Tlalnepantla, State of México 54090, México; Paul G. Allen School for Global Health, College of Veterinary Medicine, Washington State University, Pullman, Washington, USA; Department of Microbiology and Molecular Genetics, Larner College of Medicine, University of Vermont, Burlington, VT 05405, USA; Department of Microbiology & Immunology, University of Illinois Chicago, Chicago, IL USA; Chiba University-UC San Diego Center for Mucosal Immunology, Allergy, and Vaccines (CU-UCSD cMAV), La Jolla, CA 92093, USA

## Abstract

Non-typhoidal *Salmonella* serovars, such as *Salmonella enterica* serovar Typhimurium (STm), are a leading cause of inflammatory diarrhea in otherwise healthy individuals. Among children, the elderly, and immunocompromised individuals, STm can spread to systemic sites and cause potentially lethal bacteremia. Phagocytic cells and the immune complement system are pivotal to preventing the dissemination of STm. PgtE, an STm outer membrane protease, has been previously described to cleave over a dozen mammalian protein substrates *in vitro*, including complement protein C3. However, these activities have mostly been observed with mutant, avirulent strains with a truncated O-antigen that renders bacteria sensitive to complement killing. Here, we report that virulent STm utilizes PgtE to evade complement-mediated killing *in vivo*. The wild-type pathogen increases *pgtE* expression and PgtE proteolytic function within macrophages and in macrophage-like *in vitro* growth conditions, concomitant with physiologic O-antigen shortening in these environments. Furthermore, we found that wild-type STm’s resistance to complement-mediated serum and neutrophil killing is PgtE-dependent. We propose that PgtE promotes the systemic spread of STm by acting as a second line of defense against complement when STm escapes from a macrophage.

## INTRODUCTION

Infections with non-typhoidal *Salmonella* (NTS) are among the leading causes of gastrointestinal disease worldwide (1). Clinically, NTS infection presents with inflammatory diarrhea (2), characterized by localized gastrointestinal inflammation and neutrophil influx in the intestinal mucosa (3). In healthy individuals, NTS infection remains localized to the gut (2). However, approximately 5% of patients infected with NTS develop bacteremia, a serious and potentially fatal complication (2). Children and the elderly are at risk for developing bacteremia (4), and additional risk factors include leukemia, chemotherapy, and HIV infection prior to the advent of antiretroviral therapy (5–8). In recent years, invasive non-typhoidal *Salmonella* (iNTS) strains have emerged as a prominent cause of bloodstream infection in sub-Saharan Africa (9), with serovars Typhimurium (STm) and Enteritidis implicated in 91% of iNTS cases (10). Important risk factors for iNTS disease in Africa are HIV infection, malaria, and malnutrition (9). Furthermore, complicated iNTS infections present a challenge for antibiotic treatment due to increased multidrug resistance (2, 11). It is thus imperative to elucidate mechanisms by which STm can evade host immune defenses to cause bacteremia.

Neutrophils are thought to play a crucial role in preventing NTS bacteremia through limiting dissemination of the pathogen from the mucosa to systemic sites. Neutropenia in patients with HIV (7) or cancer (6), as well as defective production of reactive oxygen species (ROS) in patients with chronic granulomatous disease (12), heightens the risk of NTS bacteremia. Experiments in mice, largely conducted with STm, corroborate these clinical observations, as neutrophil depletion leads to increased pathogen dissemination (13). Even with a fully functional immune system, macrophages are less effective at killing STm due to the pathogen’s numerous strategies for survival and replication within these cells. Within the macrophage phagosome, STm uses the two-component regulatory system PhoPQ to sense acidification, Mg^2+^-limiting conditions, and cationic antimicrobial peptides, which together induce the expression of *Salmonella* Pathogenicity Island 2 (SPI2) effector genes (14–18). The SPI2-encoded type-3 secretion system delivers a plethora of effector proteins that prevent the fusion of the phagosome with lysosomes, allowing STm to persist in *Salmonella*-containing vacuoles (SCVs) within macrophages (19–21).

Protected inside the macrophage compartment, STm can spread to the liver, spleen, and blood while evading extracellular host defenses (22–25). In the extracellular environment, *Salmonella* is more vulnerable to complement opsonization, which contributes to host protection during bacteremia (26, 27) by mechanisms that are not completely elucidated. Long O-antigen chains of lipopolysaccharide on *Salmonella* play a crucial role in steric inhibition of complement, reducing effective membrane attack complex (MAC) formation. Consequently, STm lacking O-antigen (rough mutants) are susceptible to serum complement killing (28) and are avirulent (29, 30). Resistance to complement is also mediated by the outer membrane proteins TraT and Rck (31–33). A third outer membrane protein, PgtE, is a promiscuous protease described to cleave a dozen different substrates *in vitro* (34–39), including complement-associated proteins. Increased expression of *pgtE* has also been proposed to promote survival and dissemination of iNTS (39). Nevertheless, it is unknown whether the cleavage of complement proteins promotes STm virulence *in vivo*.

All previous studies investigating PgtE function *in vitro* used rough mutants, because the long O-antigen in wild-type strains sterically inhibits PgtE function (35–37, 39, 40). We thus sought to unravel the *in vivo* role of PgtE in wild-type, virulent strains with an intact O-antigen (smooth strains). Here we show that an STm *pgtE* mutant is attenuated in wild-type mice, but is rescued in complement-deficient mice. Mechanistically, we found that wild-type STm cleaves complement C3 in a PgtE-dependent manner when inside macrophages or cultured in media mimicking the SCV, environments where STm expresses a shorter O-antigen. Unexpectedly, however, PgtE-mediated disruption of complement did not promote STm survival in macrophages, but rather enhanced serum resistance and evasion of neutrophil killing, thereby contributing to bacteremia.

## MATERIALS AND METHODS

### Bacterial strains and culture conditions

Bacterial strains used in this study are listed in *Supplementary Table 1*. Plasmids used in this study are listed in *Supplementary Table 2.* Most of the *in vitro* and all of the *in vivo* work was performed with *Salmonella enterica* serovar Typhimurium (STm) strain IR715, a fully virulent, nalidixic acid-resistant derivative of strain ATCC 14028s, as well as an isogenic *pgtE* mutant of IR715. For some *in vitro* experiments, we employed the *Salmonella enterica* serovar Typhimurium sequence type ST313 strain D23580 and its isogenic *pgtE* mutant (39).

IR715 and D23580 strains were cultured on LB agar plates that were supplemented with 50 µg/ml nalidixic acid or 30 µg/ml chloramphenicol, respectively. IR715 and *E*. *coli* XL1-Blue strains transformed with a low-copy plasmid (pWSK29) encoding wild-type *pgtE* (pPgtE) or a *pgtE* inactive mutant (pPgtE*-*D206A) were grown on LB agar plates supplemented with 100 µg/ml carbenicillin. For each inoculum, three colonies were cultured overnight in 5ml of medium without antibiotic selection. All bacteria were cultured with shaking/rolling, unless otherwise stated. For animal infections, all strains were cultured in L broth (LB; 10 g/L tryptone, 5 g/L yeast extract, 10 g/L NaCl) aerobically at 37 °C, overnight. For *in vitro* experiments, strains were cultured in either LB or SPI2-inducing phosphate-carbon-nitrogen (PCN) liquid media supplemented with low magnesium (InSPI2 LowMg^2+^) (41), aerobically at 37 °C, overnight.

### Generation of bacterial mutants

Primers used in this study are listed in *Supplementary Table 3*. The STm *pgtE* mutant was constructed by allelic exchange with the plasmid pGP704 containing a tetracycline resistance cassette flanked by 1 kb regions upstream and downstream of the *pgtE* gene. Primers were used to PCR amplify 1kb upstream (left border, LB) and downstream (right border, RB) of the *pgtE* gene. The resulting products were fused in a fusion PCR and cloned into vector pCR-Blunt II-TOPO (Invitrogen). The resulting plasmid, pCRII*::pgtE*-LBRB, was sequenced and subsequently cut with SalI and EcoRV. The *pgtE*-LBRB fragment was gel purified and cloned into the SalI and EcoRV digested vector pGP704 and transformed into *E. coli* CC118 **λ***_pir_*. The resulting plasmid, pGP704*::pgtE*-LBRB, was cut with XbaI, and an NheI-digested tetracycline resistance cassette (*tetRA*) from pSPN23 was ligated into the plasmid and again transformed into CC118 **λ***_pir_*. The resulting plasmid, pGP704*::pgtE*-LBRB*::tetRA*, was transformed into *E. coli* S17-1 **λ***_pir_*, then the strain was conjugated with STm IR715, generating strain IR715 Δ*pgtE* via after selecting and screening for double-crossover events from homologous recombination. The integration of the resistance cassette and the deletion of the *pgtE* gene were confirmed by Southern blot using a probe for the 1kb region upstream of *pgtE*, and the North2South Chemiluminescent Hybridization and Detection kit (Thermo Fisher). D23580 Δ*pgtE* was constructed by transducing the *pgtE* deletion from IR715 to D23580 with P22 HT105/1 *int-201*.

For constitutive expression of the mCherry fluorescent protein, STm strains were transduced with a P22 lysate derived from STm SL1344 *glmS*::*Ptrc-mCherryST::Cm* (42), followed by removal of the Cm^R^ cassette using pCP20 (43).

For clean insertion of the FLAG sequence at the C-terminus of the chromosomal *pgtE* gene, primers for Gibson assembly were designed with the NEBuilder Assembly Tool (https://nebuilder.neb.com/#!/). FLAG_Downstream_Fwd and FLAG_Upstream_Rev primers respectively carried the FLAG sequence extension (GAC TAC AAG GAC GAC GAT GAC AAG) and the reverse complement of the FLAG sequence. Chromosomal IR715 DNA was PCR-amplified with the primer pairs of FLAG_Upstream_Fwd and FLAG_Upstream_Rev, and FLAG_Downstream_Fwd and FLAG_Downstream_Rev by PCR with High-Fidelity PCR Master Mix with HF buffer (New England Biolabs #M0531S) per manufacturer’s instructions. The plasmid pRDH10 was digested with the restriction enzymes NruI (New England Biolabs #R3192S) and SphI-HF (New England Biolabs #R3182S) per manufacturer’s instructions. All three products were then run on a 1% agarose gel, purified with a Zymoclean Gel DNA recovery kit (Zymo Research #D4001), and assembled with NEBuilder Hifi DNA assembly master mix at a 2:1 molar ratio (New England Biolabs #E5520S) following manufacturer’s instructions.

An aliquot of 100 µL of chemically competent CC118 **λ***pir* was thawed on ice then incubated with 2 µL of Gibson assembly product on ice for 30 minutes. Cells were then incubated at 42 °C in a water bath for 45 seconds, incubated on ice for 5 minutes, diluted with 1 mL of LB, and cultured for 1 hour aerobically at 37 °C. Cells were then spread-plated on LB agar plates that were supplemented with 30 µg/ml chloramphenicol, incubated overnight at 37 °C, then screened for tetracycline resistance the following day. After confirming correct Gibson assembly via sequencing of the plasmid by Primordium Labs, chemically competent S17-1 **λ***pir* cells were transformed as above with pRDH10*::pgtE*-FLAG isolated via QIAprep Spin Miniprep kit (Qiagen #27106) from CC118 **λ***pir* pRDH10*::pgtE*-FLAG. The resulting strain was used to conjugate the plasmid to STm IR715. Following conjugation, cells were incubated on LB agar plates to screen for resistance to both nalidixic acid and chloramphenicol. Cells that had undergone plasmid integration into the chromosome (single crossover events) were then counter-selected using Nutrient Broth with 7% sucrose (*sacB* gene residing in pRDH10). Clean insertion of chromosomal *pgtE*-FLAG was confirmed by PCR with primer pair FLAG_Verification_Fwd and FLAG_Verification_Rev, followed by sequencing by Primordium.

### Complementation and reporter plasmids

To construct the PgtE complementation plasmid, the *pgtE* region was PCR-amplified from STm genomic DNA. A 300 bp region upstream of the coding sequence was amplified to include relevant regulatory elements. The PCR product was cloned into plasmid pCR-Blunt II-TOPO using the Zero Blunt TOPO PCR Cloning Kit (Invitrogen) following the manufacturer’s protocol. The product was then subcloned into the multiple cloning site of low-copy plasmid pWSK29 using XhoI and EcoRV to generate plasmid pWSK29::*pgtE* (pPgtE). A missense point mutation was introduced into pWSK29::*pgtE* using the QuikChange Site-Directed Mutagenesis Kit (Agilent) to create pWSK29::*pgtE*-D206A. Sequences were confirmed by Sanger sequencing (Eton Bioscience) or Oxford Nanopore Technology (Primordium Labs).

To construct the *pgtE* reporter plasmid, the *pgtE* promoter was amplified from STm SL1344 genomic DNA with the oligos PpgtE-XbaI-F (engineered restriction sites are underlined) and PpgtE-SmaI-R. The amplicon was digested with *Xba*I/*Sma*I and ligated into *Xba*I/*Sma*I-digested pGFPmut3.1, then the *pgtE-gfpmut3.1* cassette was excised by *Xba*I/*Apa*I digestion, and ligated into the corresponding sites of pMPM-A3ΔPlac.

### Serum and serum treatments

Normal human serum (NHS; #NHS), C3-depleted human serum (#A314), and cobra venom factor (CVF; #A150) were procured from Complement Technology. For mouse serum, blood was collected from uninfected C3^+/+^ and C3^-/-^ mice through cardiac puncture with a 25-gauge needle. Mouse serum was subsequently recovered by centrifugation of blood for 5 minutes at 10,000 x *g* using Serum Gel Polypropylene Microtubes (Sarstedt, #41.1378.005). The serum was then pooled from several mice, aliquoted, and stored at - 80 °C. Both human and mouse sera were used after thawing a maximum of one time.

### Mice

The Institutional Animal Care and Use Committee (IACUC) at UC San Diego approved all mouse experiments perfomed at the institution (protocol #S17107). The IACUC at Washington State University approved mouse bone marrow collection for the generation of bone marrow-derived macrophages (protocol #6785). Mice were housed under specific pathogen-free conditions and were provided with an irradiated 2020X Teklad diet (Envigo). Furthermore, mice were randomly grouped in cages, with a maximum of five animals per cage.

The study utilized C57BL/6 wild-type mice, *C3^-/-^* mice (44), and *Cybb*-deficient mice (The Jackson Laboratory #002365) (45). For *in vivo* experiments depleting complement with CVF, six-to-eight-week-old female C57BL/6J mice (The Jackson Laboratory) were intraperitoneally injected with 0.1ml of phosphate-buffered saline (PBS) or 12.5 (one experiment) or 25 (two experiments) µg/ml CVF one day before bacterial infection (46). For all other experiments, six-to-ten-week-old female and male mice, bred and housed at UC San Diego, were used in the experiments, with similar numbers of female and male mice in each experimental group. For experiments with *C3^-/-^* mice, we used wild-type littermate control mice from the same colony (C57BL/6 background). *Cybb*-deficient mice were bred homozygous (*Cybb^X-/X-^*females) or hemizygous (*Cybb^X-/Y^* males).

For all *in vivo* experiments, STm strains were cultured aerobically in LB at 37 °C overnight. Mice were intraperitoneally infected with 1x10^4^ colony-forming units (CFUs) of STm. Blood was collected via cardiac puncture with a 25-gauge needle and syringe pre-coated with 0.5M EDTA to prevent coagulation. Liver and spleen tissues were homogenized in PBS, and samples were plated on LB agar supplemented with 50 µg/ml nalidixic acid.

### Cell culture reagents

For cell culture media, we primarily used RPMI 1640 medium with L-glutamine and Phenol Red (Gibco #11875093). In luminol assays, we employed RPMI 1640 medium with no glutamine and no phenol red (Gibco #32404014). As indicated in the respective sections, RPMI was supplemented with the following components, depending on the experiment: heat-inactivated Fetal Bovine Serum (HI-FBS) (Gibco #A3840001), Antibiotic-Antimycotic solution (Gibco #15240062), Gentamicin (Gibco #15710064), HEPES (Gibco #15630080), EDTA (Fisher Scientific #S311-500). Dulbecco’s PBS (DPBS; Gibco #14190) was used for dislodging bone marrow-derived macrophages and for the neutrophil Enrichment Kit isolation medium.

### Bone marrow isolation and bone marrow-derived macrophage culture conditions

Murine bone marrow-derived macrophages (BMDMs) were prepared by maturing freshly isolated bone marrow cells from femurs and tibias. Bone marrow cells were isolated with a 21-gauge needle, filtered through a 70 µm filter, then subjected to Ammonium-Chloride-Potassium (ACK) lysis (150 mM NH_4_Cl, 10 mM KHCO_3_, 0.1 mM Na_2_EDTA) buffer to remove excess red blood cells. For BMDMs used in fluorescent microscopy, cells were cultured for 5 days in RPMI 1640 medium with L-glutamine supplemented with 20% supernatant from L929 cells, and 10% HI-FBS. BMDMs were then re-seeded two days prior to infection. For BMDMs used to assess *Salmonella* burden and PgtE function, cells were then cultured for 7 days in RPMI 1640 medium with L-glutamine supplemented with 30% supernatant from L929 cells, 10% HI-FBS, and 1x Antibiotic-Antimycotic in Sigma culture dishes (Z358762). 18 hours prior to infection, cold DPBS was used to dislodge the cells, and BMDMs were seeded in RPMI 1640 medium with L-glutamine supplemented with 10% HI-FBS in 24-well plates (Corning #3524) at a density of 5x10^5^ cells/well or 6-well plates at a density of 2x10^6^ cells/well (Corning #3516).

### Murine macrophage infection for bacterial enumeration

For macrophage infection experiments, STm strains were grown statically in LB media in an aerobic environment at 37 °C overnight. A concentration of 1.67x10^7^ CFU/ml of STm was incubated in 20% mouse serum (opsonized) or PBS (non-opsonized) for 30 minutes at room temperature. Subsequently, STm was diluted 1:10 in RPMI 1640 medium with L-glutamine supplemented with 10% HI-FBS for an inoculum of 2% mouse serum with 1.67x10^6^ CFU/ml STm. An aliquot of 300uL of this inoculum was added to BMDMs in a 24-well plate to reach an MOI of 1. The plate was centrifuged at 360 x *g* for 5 minutes at room temperature then transferred to a 37 °C tissue culture incubator. After 30 minutes of infection, BMDMs were washed with PBS then treated with RPMI 1640 medium with L-glutamine supplemented with 10% HI-FBS and 100 µg/ml gentamicin for 30 min before replacement with RPMI 1640 medium with L-glutamine supplemented with 10% HI-FBS and 20 µg/ml gentamicin for the remainder of the assay. BMDMs were washed with PBS then lysed with 1% Triton X-100 surfactant (EMD Millipore #EM-9400) in PBS at 30 minutes, 8 hours, and 24 hours post-infection. CFUs were enumerated by plating aliquots of serially diluted lysates onto LB agar supplemented with 50 µg/ml nalidixic acid.

### Western blot detection of PgtE-FLAG and PgtE-dependent C3 cleavage

To assess PgtE-dependent cleavage of C3 *in vitro*, strains of STm and *E. coli* XL1-Blue were cultured overnight in LB or in InSPI2 LowMg^2+^ media in an aerobic environment at 37 °C. Bacteria were then incubated with 20% normal human serum (NHS) in PBS at 1.67x10^9^ CFU/ml for 8 hours. Samples were subsequently centrifuged at 10,000 x *g* for 5 minutes, and supernatants were collected for Western blotting.

To assess PgtE-dependent cleavage of C3 by intracellular STm isolated from BMDMs, STm strains were cultured by rotating in LB media in an aerobic environment at 37 °C overnight. STm was incubated in 20% mouse serum in PBS for 30 minutes at 37 °C at a concentration of 2x10^7^ CFU/ml. STm was then diluted 1:40 in RPMI 1640 medium with L-glutamine supplemented with 10% HI-FBS, then added to BMDMs in a 6-well plate at an MOI of 10. Plates were centrifuged at 360 x *g* for 5 minutes at room temperature and then transferred to a 37 °C tissue culture incubator. After 30 minutes of infection, BMDMs were washed with PBS then treated with RPMI 1640 medium with L-glutamine supplemented with 10% HI-FBS and 100 µg/ml gentamicin for 30 min before replacement with RPMI 1640 medium with L-glutamine supplemented with 10% HI-FBS and 20 µg/ml gentamicin for 7.5 hours. Infected BMDMs were then washed with PBS and lysed with water for 10 minutes at 37 °C. Six infected wells were pooled together for each group, washed, resuspended in 100 µl of 20% NHS in PBS, then shaken at 300 rpm at 37 °C for 13 hours. Samples were then centrifuged at 10,000 x *g* for 5 minutes, and supernatants were collected for western blotting.

To assess PgtE protein production by *in vitro* cultures, STm WT and STm *pgtE*-FLAG (strain ML27) were cultured overnight in LB or in InSPI2 LowMg^2+^ media in an aerobic environment at 37 °C. 5x10^8^ CFUs were washed twice in PBS; pellets were frozen at - 80 °C for 30 minutes, then resuspended in 50 µl of lysis buffer (2% 2-Mercaptoethanol, 2% SDS, 10% glycerol, and 0.1M TrizmaHCl in water adjusted to pH 6.8). Samples were incubated at 95 °C for 20 minutes then spun down for 10 minutes at 10,000 x *g*.

For electrophoresis, samples were prepared with RunBlue LDS Sample Buffer (Expedeon #NXB31010) and 5mM dithiothreitol (Thermo Scientific #R0861). Electrophoresis was conducted using a Mini Gel Tank (Invitrogen #A25977), Novex Tris-Glycine Mini Protein Gel 4-12% (Invitrogen #XP04125BOX), WesternSure Pre-stained Chemiluminescent Protein ladder (Li-Cor #926-98000) and MES SDS Running Buffer (Invitrogen #B0002) at 90 volts for 80 minutes. Semi-dry transfer was performed with a Trans-Blot SD Semi-Dry Transfer Cell (Bio-Rad), Immun-Blot PVDF membrane (Bio-Rad #1620177), and Whatman GB003 gel blotting papers (Whatman #10427806) at 20 volts for 1 hour.

Membranes were blocked with 5% (w/v) Nonfat dry milk (LabScientific #M0841) in Tris-buffered saline with 0.1% (w/v) Tween 20 (TBST) rocking for 2 hours at room temperature. For PgtE-dependent complement cleavage, membranes were then incubated with purified anti-complement C3/C3b/iC3b/C3d antibody (BioLegend #846302 clone 1H8/C3b) diluted to 1:5,000 in 5% milk in TBST rocking overnight at 4 °C. After 5 washes with TBST, membranes were then incubated with HRP goat anti-mouse IgG (BioLegend #405306) diluted to 1:20,000 in 5% milk in TBST rocking overnight at 4 °C. For detection, membranes were washed 5 times with TBST, incubated for 10 minutes in the dark with ECL Prime Western Blotting Detection Reagents (Amersham #RPN2232), and then imaged with an Azure 300 Chemiluminescent Western Blot Imager (Azure Biosystems #AZ1300-01).

For PgtE-FLAG tag analysis, after semi-dry transfer, PVDF membranes were cut in half at the 50 kDa protein ladder mark. The bottom half of the membrane was then incubated with purified rat anti-DYKDDDDK Tag antibody (anti-FLAG tag; BioLegend #637319 clone L5) diluted to 1:5,000 in 5% milk in TBST rocking overnight at 4 °C. After 5 washes with TBST, membranes were then incubated with HRP goat anti-rat IgG (BioLegend #405405) diluted to 1:5,000 in 5% milk in TBST rocking overnight at 4 °C. The top half of the membrane was incubated with mouse anti-DnaK (*E. coli*) antibody (Enzo #ADI-SPA-880-D clone 8E2/2) diluted to 1:10,000 in 5% milk in TBST, rocking overnight at 4 °C. After 5 washes with TBST, membranes were then incubated with HRP goat anti-mouse IgG antibody (BioLegend #405306) diluted to 1:10,000 in 5% milk in TBST rocking overnight at 4 °C. For detection, membranes were washed 5 times with TBST, incubated for 10 minutes in the dark with ECL Prime Western Blotting Detection Reagents (Amersham #RPN2232), and then imaged with a GeneGnome (Synoptics).

### O-Antigen Staining

STm and *E. coli* XL1-Blue strains were cultured overnight in LB or in InSPI2 LowMg^2+^ media in an aerobic environment at 37 °C. 5x10^8^ CFU was washed twice in PBS and then resuspended in 100 µl of lysis buffer (2% 2-Mercaptoethanol, 2% SDS, 10% glycerol, and 0.1M TrizmaHCl in water adjusted to pH 6.8). Samples were incubated at 95 °C for 10 minutes and then incubated with 1.25 µl of Proteinase K (20mg/ml; Viagen #501-PK) overnight at 55 °C. Lysates were prepared for electrophoresis with Laemmli Sample Buffer (Bio-Rad #1610747) and 7.5% 2-Mercaptoethanol. Electrophoresis was conducted using a Mini Gel Tank (Invitrogen #A25977), Novex Tris-Glycine Mini Protein Gel 4-12% (Invitrogen #XP04125BOX), and MES SDS Running Buffer (Invitrogen #B0002) at 25 mA for 2 hours. O-antigen staining was then performed with Pro-Q Emerald 300 Lipopolysaccharide Gel Stain Kit (Invitrogen #P20495) following the manufacturer’s instructions. Gels were imaged with the 302 nm UV transilluminator of an Azure 200 (Azure Biosystems #AZ1200-01).

### Mouse neutrophil isolation

Fresh femur- and tibia-isolated bone marrow cells were isolated with a 21-gauge needle and filtered through a 70 µm filter. Neutrophils were isolated with the EasySep Mouse Neutrophil Enrichment Kit (Stemcell Technologies #19762) following the manufacturer’s instructions for the EasySep Magnet (Stemcell Technologies #18000). The isolation medium consisted of DPBS supplemented with 2% HI-FBS and 1 mM EDTA.

### Neutrophil killing assay

Murine bone marrow neutrophils were resuspended in RPMI 1640 medium with L-glutamine supplemented with 10% HI-FBS and 1mM HEPES, then plated at 5x10^5^ cells/well in a 96-well round bottom cell culture plate (Costar #3799). Neutrophils were incubated in a 37 °C tissue culture incubator for 30 minutes prior to infection.

STm strains were cultured overnight in LB or in InSPI2 LowMg^2+^ media in an aerobic environment at 37 °C. A concentration of 5x10^8^ CFU/ml of STm was incubated in 20% mouse serum from C3^+/+^ and C3^-/-^ mice (opsonized) or PBS (non-opsonized) for 30 minutes at room temperature. STm was then diluted 1:10 in RPMI 1640 medium with L-glutamine supplemented with 10% HI-FBS and 1mM HEPES, resulting in an inoculum of 5x10^7^ CFU/ml STm with 2% mouse serum. Subsequently, 100 µl of inoculum was added to wells with 100 µl of medium or 100 µl of 5x10^5^ neutrophils for an MOI of 10. After 2.5 hours in a 37 °C tissue culture incubator, 100 µl of 2% Triton X-100 surfactant in PBS was added to 100 µl of culture. CFUs were enumerated by plating aliquots of serially diluted lysates onto LB agar supplemented with 50 µg/ml nalidixic acid.

### Luminol Assay

STm strains were grown aerobically overnight at 37 °C, then sub-cultured in LB (1:100 dilution) or in InSPI2 LowMg^2+^ media (1:10 dilution) and grown aerobically at 37 °C for 3 hours. A concentration of 1x10^8^ CFU/ml of STm was then incubated in 20% mouse serum from C3^+/+^ and C3^-/-^ mice for 30 minutes at room temperature. Murine bone marrow neutrophils were resuspended in RPMI 1640 medium with no glutamine and no phenol red supplemented with 2% HI-FBS and 1mM Luminol (Millipore Sigma #123072-2.5g) at 1.11x10^6^ neutrophils/ml. 90 µl of 1.11x10^6^ neutrophils/ml were added to a white opaque 96-well microplate (OptiPlate-96; Revvity #6005290). The plate was sealed with a Breathe-Easy sealing membrane (Diversified Biotek #BEM-1), and baseline luminescence was measured with a Synergy HTX Multi-Mode Microplate Reader (Agilent, formerly BioTek) at 37 °C. An aliquot of 10 µl of opsonized STm was then quickly added to each well for a final concentration of 10^6^ neutrophils/ml, an MOI of 10, and a final concentration of 2% mouse serum, then resealed with Breathe-Easy sealing membrane. Luminescence was recorded every 2 minutes for 120 minutes.

### Fluorescence Microscopy

Infected macrophages were fixed in 2.5% (w/v) paraformaldehyde at 37 °C for 10 min then washed three times in PBS. Monolayers were permeabilized in 10% (v/v) normal goat serum (Life Technologies), 0.2% (w/v) saponin in PBS for 20 min at room temperature, incubated with primary antibodies for 45 min at room temperature, washed three times with 0.2% (w/v) saponin in PBS, then incubated with secondary antibodies for 45 min at room temperature. Coverslips were washed in PBS, incubated with Hoechst 33342 (ThermoFisher Scientific) for 1 min to stain DNA, and then mounted onto glass slides in Mowiol (Calbiochem). Samples were viewed with a Leica DM4000 epifluorescence upright microscope for quantitative analysis or a Leica SP8 confocal laser-scanning microscope for image acquisition. Samples were blinded during the experiment. Representative confocal micrographs of 1024x1024 pixels were acquired and assembled using Adobe Photoshop CS6.

### Statistical analysis of data

The experiments were not randomized. No statistical methods were used to predetermine the sample size. Prism 10 software (GraphPad) was used for statistical analysis. For *in vivo* experiments, outliers found by ROUT outlier analysis Q= 1% are removed. Data were analyzed by Kruskal-Wallis test (non-parametric, no pairing) followed by Dunn’s multiple comparison test. Serum killing assays were analyzed with a Two-way ANOVA followed by Sidak multiple comparison test. Neutrophil killing assays were analyzed with a One-way ANOVA Kruskal-Wallis test followed by Dunn’s comparison test. For luminol assays, Two-way ANOVA analysis was performed; the source of variation for significance is the Time x Column Factor.

## RESULTS

### PgtE promotes immune complement resistance *in vivo*

Prior studies identified a potential role for PgtE in promoting STm colonization in mice and chickens (37, 39, 47) and described several potential proteolytic targets *in vitro*, including complement factor B, complement factor H, C3, C3b, C4b, and C5 (36, 38, 39).

All three immune complement pathways converge at C3 (48). To elucidate whether PgtE enables STm to evade immune complement *in vivo*, we infected *C3^-/-^* mice and their *C3^+/+^* littermates intraperitoneally with STm WT (strain IR715, a fully virulent Nal^R^ derivative of ATCC 14028s) or an isogenic Δ*pgtE* mutant (**Fig. 1A-E**). After 24 hours, we assessed bacterial burden in the blood (**Fig. 1B**), liver (**Fig. 1C**), and spleen (**Fig. 1D**). The Δ*pgtE* mutant was recovered at significantly lower levels than STm WT in the blood, but was fully rescued in *C3^-/-^* mice (**Fig. 1B**). Similar differences between STm WT and the Δ*pgtE* mutant were observed in the liver and spleen of *C3^+/+^* mice, although they did not reach statistical significance. In all cases, while STm WT equally infected *C3^+/+^*and *C3^-/-^* mice, the Δ*pgtE* mutant was recovered at much higher levels in the spleen and liver of *C3^-/-^* mice when compared to *C3^+/+^* littermates (**Fig. 1C, D**). Furthermore, *C3^-/-^*mice infected with the Δ*pgtE* mutant exhibited significantly higher weight loss than the infected *C3^+/+^* mice (**Fig. 1E**). Thus, PgtE enables STm to evade immune complement defense *in vivo*, particularly in the blood.

**Figure 1.**
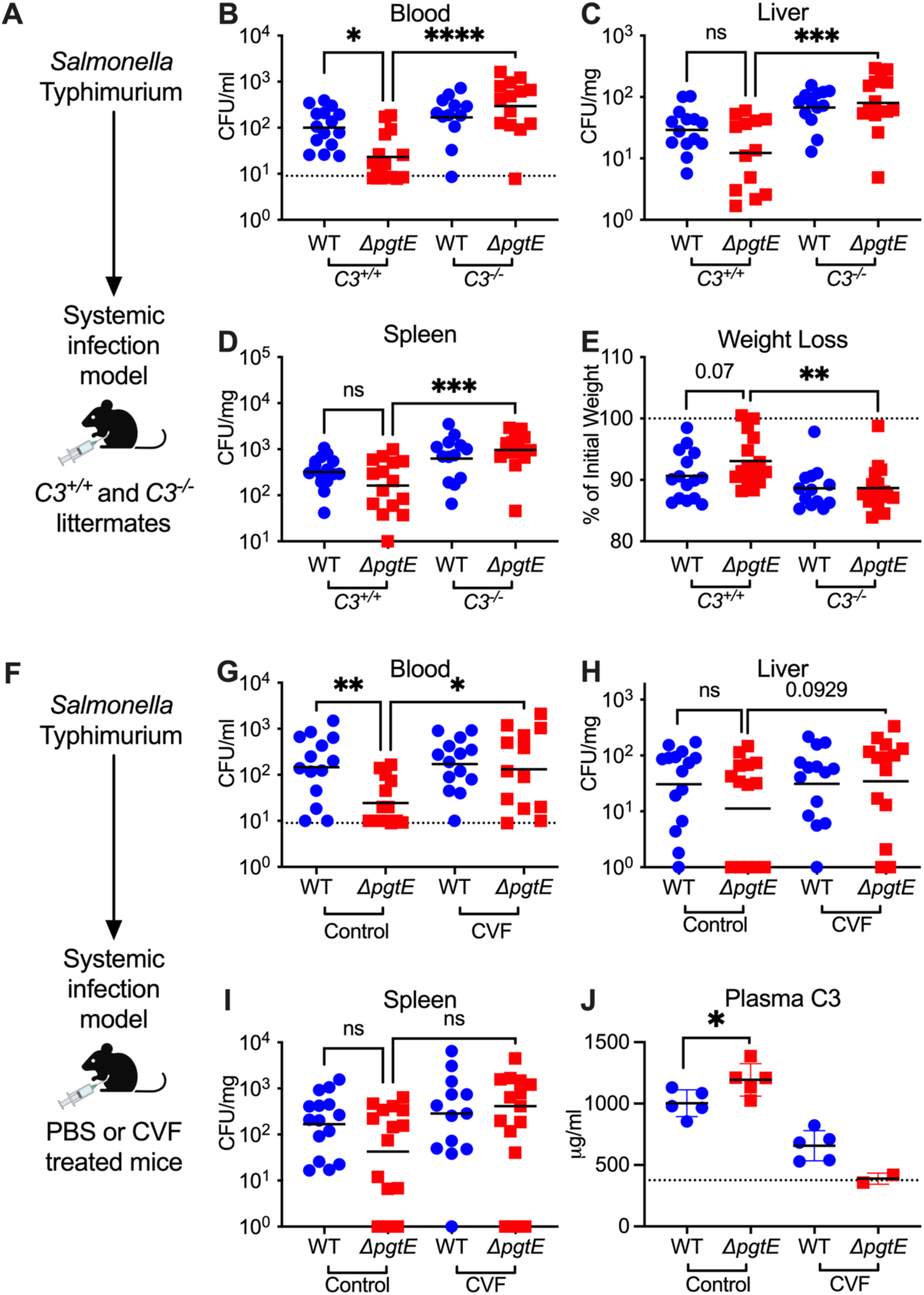
PgtE promotes smooth STm survival *in vivo* by evading complement C3. (**A-E**) 6-10-week-old *C3^+/+^*and *C3^-/-^* littermates were infected intraperitoneally (IP) with 10^4^ CFU wild-type (WT) or isogenic PgtE-deficient (*ΔpgtE*) *Salmonella* strain IR715. Mice were euthanized 24 hours after infection and bacterial burden in the (**B**) blood, (**C**) liver, and (**D**) spleen were quantified. (**E**) Weight loss = (weight at 24 hours / weight at time of infection)*100%. (**F-J**) 6-8-week-old C57B6/J mice were IP-injected with PBS (Control) or Cobra Venom Factor (CVF). 24 hours after treatment, mice were infected IP with 10^4^ CFU of either IR715 WT or IR715 *ΔpgtE*. Mice were euthanized 24 hours after infection and bacterial burden was assessed in the (**G**) blood, (**H**) liver, and (**I**) spleen. (**J**) Concentration of complement C3 in plasma measured by ELISA: dotted line represents average from 3 uninfected control mice. (**B, G**) Dotted line represents the limit of detection of STm CFU in blood. (**B-E**) N = 16-17 per group pooled from 6 independent experiments. (**G-I**) N = 15 per group pooled from 3 independent experiments. (**J**) ELISA from 1 representative experiment. (**B-E, G-I**) Outliers found by ROUT outlier analysis Q= 1% are removed. Data were analyzed by Kruskal-Wallis test (non-parametric, non-paired) followed by Dunn’s multiple comparison test. Adjusted p values from Dunn’s multiple comparison test: * p < 0.05. ** p < 0.01. *** p < 0.001. ns = not significant. Symbols represent data from individual mice. Bars represent the (**B-D, G-I**) geometric means or (**E, J**) mean.

We further investigated PgtE-dependent evasion of complement by infecting mice treated with cobra venom factor (CVF), a C3 convertase homolog which depletes complement (46) (**Fig. 1F-J**). Mice treated with PBS (control) or CVF for 24 hours were infected intraperitoneally with STm WT or the Δ*pgtE* mutant (**Fig. 1F**), and bacterial burden was assessed in the blood (**Fig. 1G**), liver (**Fig. 1H**), and spleen (**Fig. 1I**) at 24 hours. Similar to *C3*^+/+^ mice, the Δ*pgtE* mutant was recovered at significantly lower levels than STm WT in the blood of control-treated mice but was rescued in CVF-treated mice (**Fig. 1G**). No significant differences were observed in the liver (**Fig. 1H**) and spleen (**Fig. 1I**). To confirm that CVF treatment effectively depleted complement C3, we determined serum C3 concentration by ELISA. As expected, mice treated with CVF had reduced serum C3 compared to control-treated mice (**Fig. 1J**). Within the control-treated group, mice infected with STm WT had significantly less serum C3 compared to mice infected with the Δ*pgtE* mutant (**Fig. 1J**), suggesting that PgtE reduced serum C3 concentrations. Thus, PgtE enables STm to defend against immune complement *in vivo*.

### Wild-type STm cleaves complement C3 in a PgtE-dependent manner when grown in conditions that mimic the phagosome or grown in macrophages

Previous *in vitro* studies used strains with a defective O-antigen, and thus were avirulent, to show PgtE-dependent cleavage of immune complement (36, 38, 39). As we identified a potential role for PgtE in cleaving C3 *in vivo*, we hypothesized that PgtE acts by a different mechanism in fully virulent STm.

Transcriptome analysis has revealed that STm increases *pgtE* expression in infected murine macrophages (49), indicating that PgtE may function in these cells. To elucidate the time course of *pgtE* expression, we infected bone marrow-derived macrophages (BMDMs) with an STm strain carrying a chromosomally encoded P*trc*::*mCherry*, for constitutive expression of mCherry fluorescent protein and a plasmid encoding a P*pgtE*::*gfp* transcriptional reporter fusion (**Fig. 2A**). Monitoring GFP fluorescence over time by fluorescence microscopy revealed that 4.3% and 80% of bacteria were GFP-positive at 30 minutes and 8 hours post-infection, respectively (**Fig. 2B**). These results indicated a temporal induction of *pgtE* expression following STm infection of BMDMs.

**Figure 2.**
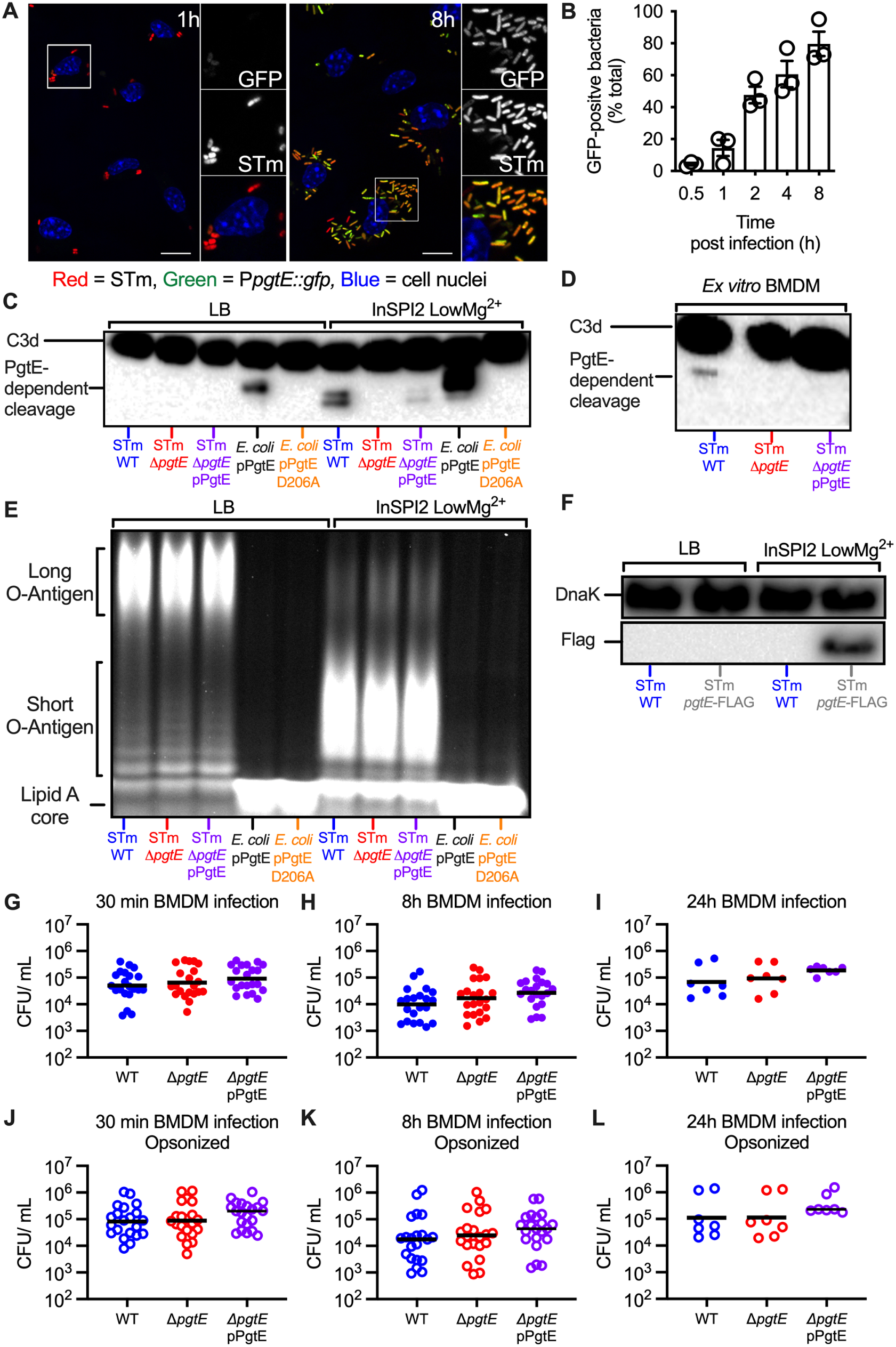
PgtE expression and function are increased in macrophages but do not increase smooth STm survival in macrophages. (**A, B**) Temporal and spatial distribution of PgtE-positive STm inside BMDMs. (**A**) BMDMs were infected with mCherry-STm carrying a plasmid encoding for a P*pgtE*::*gfp* transcriptional reporter fusion. Representative confocal microscopy images from 1 h and 8 h post-infection are displayed. GFP-positive bacteria (green), *Salmonella* (red), and the cell nuclei (DAPI; blue) are shown. Inset panels show 2x enlarged regions; scale bars are 10 μm. (**B**) Kinetics of intracellular *pgtE* expression in BMDMs. The number of GFP-positive bacteria at each timepoint was scored by fluorescence microscopy and reported as a percentage of total (red) bacteria (n = 3 experiments). (**C, E**) Smooth STm IR715 wild-type (WT), isogenic PgtE-deficient (*ΔpgtE*), and *ΔpgtE* complemented in trans (*ΔpgtE* pPgtE) or rough *E. coli* with a pWSK29 plasmid containing a functional *pgtE* gene (pPgtE) or a *pgtE* gene with a single point mutation PgtE (pPgtE D206A) were cultured overnight in (**Left**) LB or (**Right**) InSPI2 LowMg^2+^ minimal media. (**D**) Alternatively, STm was isolated from BMDMs 8 hours after infection. STm and *E. coli* were then incubated with normal human serum for (**C**) 8 hours or (**D**) 13 hours. PgtE-dependent complement cleavage in supernatants was assessed by western blot analysis with anti-complement C3/C3b/iC3b/C3d antibody. (**E**) Alternatively, after overnight culture, STm and *E. coli* were lysed, run on a 4-12% Tris-Glycine gel, and stained with Pro-Q Emerald 300 Lipopolysaccharide Gel Stain Kit to assess O-antigen chain length. (**F**) Western blot analysis of STm WT or STm *pgtE*-FLAG cultured overnight in LB or InSPI2 LowMg^2+^ minimal media. The bottom half of the membrane was stained with anti-FLAG tag antibody. The top half of the membrane was stained with anti-DnaK as a loading control. (**G-L**) BMDMs were infected at an MOI = 1 with IR715 WT, *ΔpgtE*, and *ΔpgtE* pPgtE that were either (**G-I**) not opsonized or (**J-L**) opsonized with normal mouse serum. (**G, J**) 30 minutes after infection, BMDM were lysed with 1% Triton-X 100 and STm CFUs were enumerated. Alternatively, BMDM were incubated with 100 µg/mL gentamicin for 30 minutes, followed by (**H, K**) 7 hours or (**I, L**) 23 hours with 20 µg/mL gentamicin then lysed with 1% Triton-X 100. (**G-L**) N = 21 or 7 from 10 or 3 independent experiments. Symbols represent data from BMDMs from individual mice, bars represent the geometric means.

The phagosome’s environment can be modeled *in vitro* using minimal phosphate-carbon-nitrogen (PCN) media supplemented with low magnesium. This medium induces SPI2 expression and is thus referred to as “InSPI2 LowMg^2+^”. In alignment with the macrophage results, *pgtE* expression is also increased in this medium (41). We thus investigated whether PgtE activity *in vitro* was dependent on culture conditions. We grew the following strains in standard LB or in InSPI2 LowMg^2+^ media: STm WT, the Δ*pgtE* mutant, and the Δ*pgtE* mutant complemented with a plasmid encoding *pgtE* (STm Δ*pgtE* pPgtE). As controls, we used an O-antigen-deficient *E. coli* strain expressing either functional *pgtE* (*E. coli* pPgtE) or nonfunctional *pgtE* with a missense point mutation (*E. coli* pPgtE D206A). Each culture was then incubated with normal human serum (NHS), which contains complement, to investigate C3 cleavage by Western blot.

In line with previous studies (35, 36, 40), STm WT grown in LB was unable to cleave C3 in a PgtE-dependent manner (**Fig. 2C**, **Left**). The O-antigen-deficient *E. coli* cleaved C3 when expressing functional PgtE, consistent with the hypothesis that long O-antigen sterically inhibits PgtE function (**Fig. 2C**, **Left**). Strikingly, however, STm WT cultured in InSPI2 LowMg^2+^ media cleaved C3 in a PgtE-dependent manner, as shown by two C3 cleavage products that were absent from sera incubated with STm Δ*pgtE* (**Fig. 2C, Right**). Genetic complementation *in trans* recovered PgtE-dependent C3 cleavage, albeit to a lesser extent than STm WT.

As InSPI2 LowMg^2+^ media models the intraphagosomal environment, we next investigated whether STm WT could cleave C3 when grown inside macrophages. We infected BMDMs with STm strains (WT, the Δ*pgtE* mutant, and the complemented strain) for 8 hours, then lysed the infected cells to retrieve STm. Bacteria isolated from macrophages were then incubated with NHS to detect their ability to cleave C3. We detected a C3 fragment in serum incubated with STm WT isolated from macrophages, but not in serum incubated with the Δ*pgtE* mutant (**Fig. 2D**). In this experimental setting, genetic complementation did not restore detectable PgtE-dependent C3 cleavage. Comparing these results with those generated with STm cultured in InSPI2 LowMg^2+^ media (**Fig. 2C, Right**), where also one additional fragment was detected, we speculate that this discrepancy is attributable to the technical limitation of isolating substantially fewer STm from infected BMDMs than from overnight cultures. Nevertheless, our results demonstrate that PgtE is functional in STm with an intact O-antigen depending on the growth conditions, enabling the pathogen to cleave C3 when cultured in InSPI2 LowMg^2+^ media or when isolated from macrophages.

### Growth conditions that model the phagosome’s environment increase PgtE expression and decrease O-antigen length

PgtE activity can be observed *in vitro* among strains with an intact O-antigen as long as they are cultured in media that mimics the intraphagosomal environment. As avirulent mutants lacking an O-antigen have previously been shown to exhibit PgtE function, and as *in vitro* culture conditions and growth in macrophages can alter O-antigen length in wild-type strains (50, 51), we sought to determine whether the O-antigen length of our virulent, smooth strains was being altered by these growth conditions. To this end, we extracted and stained the O-antigen from STm strains cultured in LB or in InSPI2 LowMg^2+^media. All STm strains cultured in InSPI2 LowMg^2+^ media had shorter O-antigen compared to STm cultured in LB (**Fig. 2E**). As expected, the rough *E. coli* strain that we used to express PgtE lacked O-antigen polysaccharides. Consistent with the observation that steric hindrance conferred by the presence of an O-antigen impacts PgtE function, PgtE activity was greatest when the protease was expressed by the rough *E. coli* strain (**Fig. 2C**). By contrast, although the shorter O-antigen detected in smooth STm strains cultured in InSPI2 LowMg^2+^ media likely enabled PgtE’s ability to function at all, the intermediary PgtE activity observed is likely the consequence of lingering steric hindrance conferred by the still present, albeit shorter, O-antigen. Nevertheless, these results are consistent with the idea that the shorter O-antigen induced by growth in InSPI2 LowMg^2+^ media enables complement C3 cleavage by PgtE (**Fig. 2C, E**).

The absence of PgtE activity when wild-type STm is cultured in LB could be due to a lack of PgtE expression or it could be solely explained by the steric hindrance caused by the long O-antigen. To assess whether PgtE is expressed in LB, we constructed an STm strain with a chromosomal *pgtE* allele harboring a FLAG tag at the C-terminus (STm *pgtE*-FLAG). We found that the FLAG tag was detectable when STm *pgtE*-FLAG was cultured in InSPI2 LowMg^2+^ medium, but not in LB (**Fig. 2F**). As expected, no FLAG tag was detected in STm WT in either condition. Thus, growth in InSPI2 LowMg^2+^ media has a two-pronged effect: 1) increasing PgtE expression; 2) shortening O-antigen length, which enables PgtE function and cleavage of complement C3.

### PgtE appears dispensable for STm survival in primary macrophages under tested conditions

Our findings suggest a role for PgtE to enable *Salmonella* survival inside of macrophages. Even though complement is generally known to opsonize and lyse pathogens in extracellular spaces, recent studies have identified a role for complement in intracellular compartments (52–54). We thus tested whether PgtE disrupts intracellular C3 signaling and promotes STm survival within macrophages by infecting BMDMs with STm WT, the Δ*pgtE* mutant, or the complemented Δ*pgtE* mutant. The strains were either nonopsonized (**Fig. 2G-I**) or opsonized with normal mouse serum (**Fig. 2J-L**). We recovered a similar number of each STm strain at each of the time points analyzed, from 30 minutes post-infection (when *pgtE* is not highly expressed; **Fig. 2A, B**) to 8 hours (high *pgtE* induction) and even 24 hours post-infection, in both the non-opsonized and the opsonized groups (**Fig. 2G-I** and **Fig. 2 J-L**). As such, PgtE did not enhance STm survival in BMDMs in these conditions, even though it is highly produced and cleaves C3 in these cells.

### PgtE increases STm serum resistance

To determine whether PgtE promotes STm resistance to serum killing, we cultured STm WT, the Δ*pgtE* mutant, and the complemented Δ*pgtE* mutant in either LB or InSPI2 LowMg^2+^ media and exposed them to 20% normal human serum (NHS). When STm was cultured overnight in LB (**Fig. 3A**), all strains showed similar survival. However, when STm was cultured overnight in InSPI2 LowMg^2+^ media, STm WT survived significantly more than the Δ*pgtE* mutant, with the complemented strain showing an intermediate phenotype (**Fig. 3B**). To test whether the differences in serum resistance were dependent on PgtE-mediated C3 cleavage, the strains were incubated with C3-depleted human serum after overnight culture in InSPI2 LowMg^2+^ media. In the absence of C3, serum survival of the PgtE mutant was fully restored, and no difference in survival was detected between the three strains (**Fig. 3C**). Thus, PgtE enhanced STm serum survival by inhibiting the function of complement.

**Figure 3.**
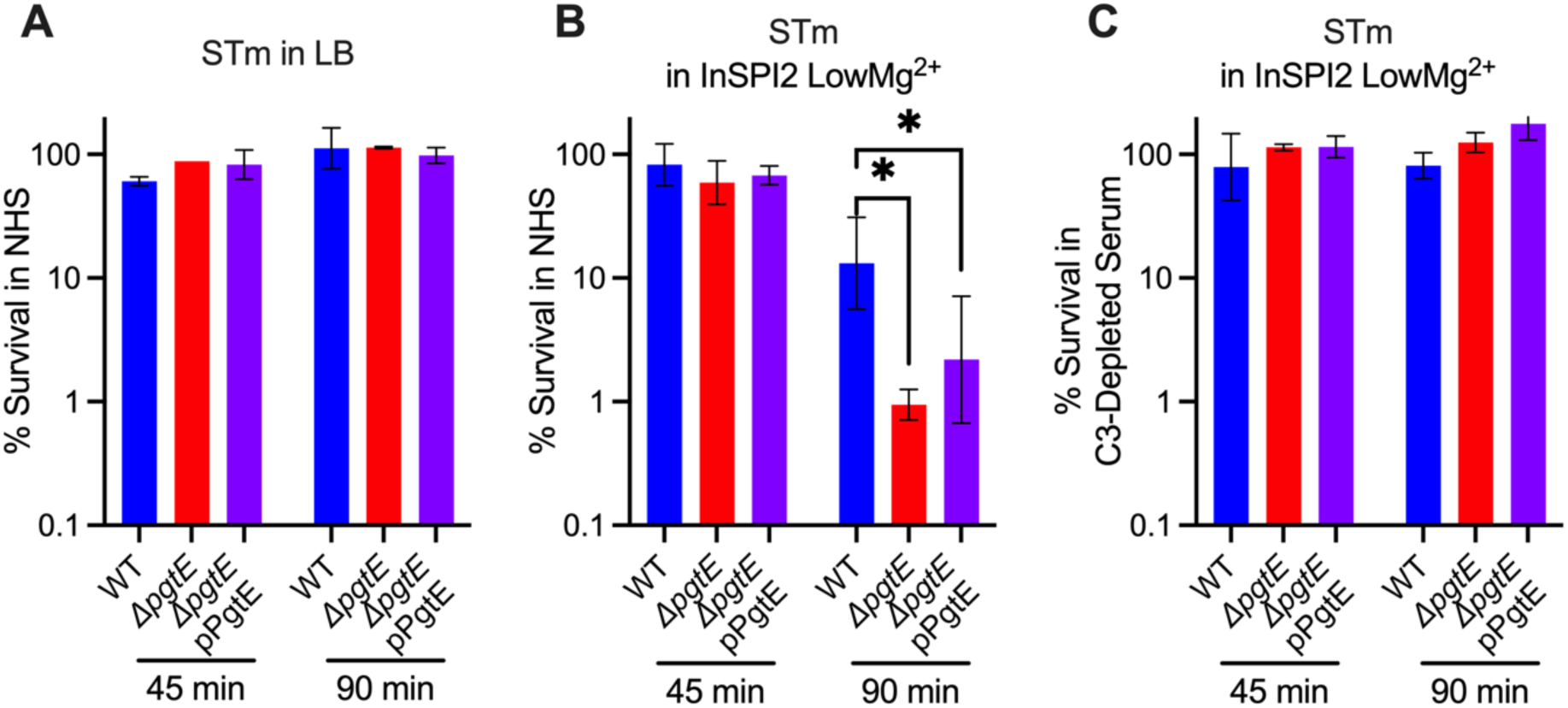
PgtE promotes survival of smooth, virulent STm in serum. (**A-C**) Serum killing assays were performed with smooth STm IR715 wild-type (WT), isogenic PgtE-deficient (*ΔpgtE*), and *ΔpgtE* complemented *in trans* (*ΔpgtE* pPgtE). Strains were cultured overnight (**A**) in LB or (**B**, **C**) in InSPI2 LowMg^2+^ minimal media. STm at 10^6^ CFU/mL was then incubated with (**A, B**) 20% normal human serum (NHS) or (**C**) 20% C3-depleted human serum at 37 °C shaking at 300 rpm. CFU were enumerated at 0 minutes, 45 minutes, and 90 minutes. % survival = (CFU at 45 minutes or 90 minutes / CFU at 0 minutes)*100%. (**A, C**) n = 2, (**B**) n = 6 from 2-3 independent experiments. Bar and error represent geometric mean and standard deviation. Data were analyzed by 2-way ANOVA followed by Sidak multiple comparison test. Adjusted p values from Sidak multiple comparison test: * p < 0.05.

Many iNTS isolates display increased expression of *pgtE* (39). We next tested if PgtE played a similar role in increasing serum survival of iNTS sequence type ST313, a predominant etiologic agent of iNTS disease (55). Similar to what we observed with the ATCC 14028s strain IR715 (sequence type ST19), no significant difference in serum survival was seen between the ST313 strain D23580 wild-type and an isogenic Δ*pgtE* mutant when the strains were cultured overnight in LB (**Fig. 4A**). However, when cultured overnight in InSPI2 LowMg^2+^ media, D23580 WT survived significantly better than the isogenic Δ*pgtE* mutant in normal human serum (**Fig. 4B**) but not in C3-depleted human serum (**Fig. 4C**). Both D23580 WT and Δ*pgtE* strains exhibited shortened O-antigen chains when cultured overnight in InSPI2 LowMg^2+^ media compared to growth in LB (**Fig. 4D**), whereas only WT was able to cleave C3 (**Fig. 4E**). Thus, akin to the results with ST19 strains (**Fig. 2C, 3**), when an ST313 strain is cultured in media mimicking the SCV, PgtE-dependent inhibition of complement results in elevated serum survival (**Fig. 4**).

**Figure 4.**
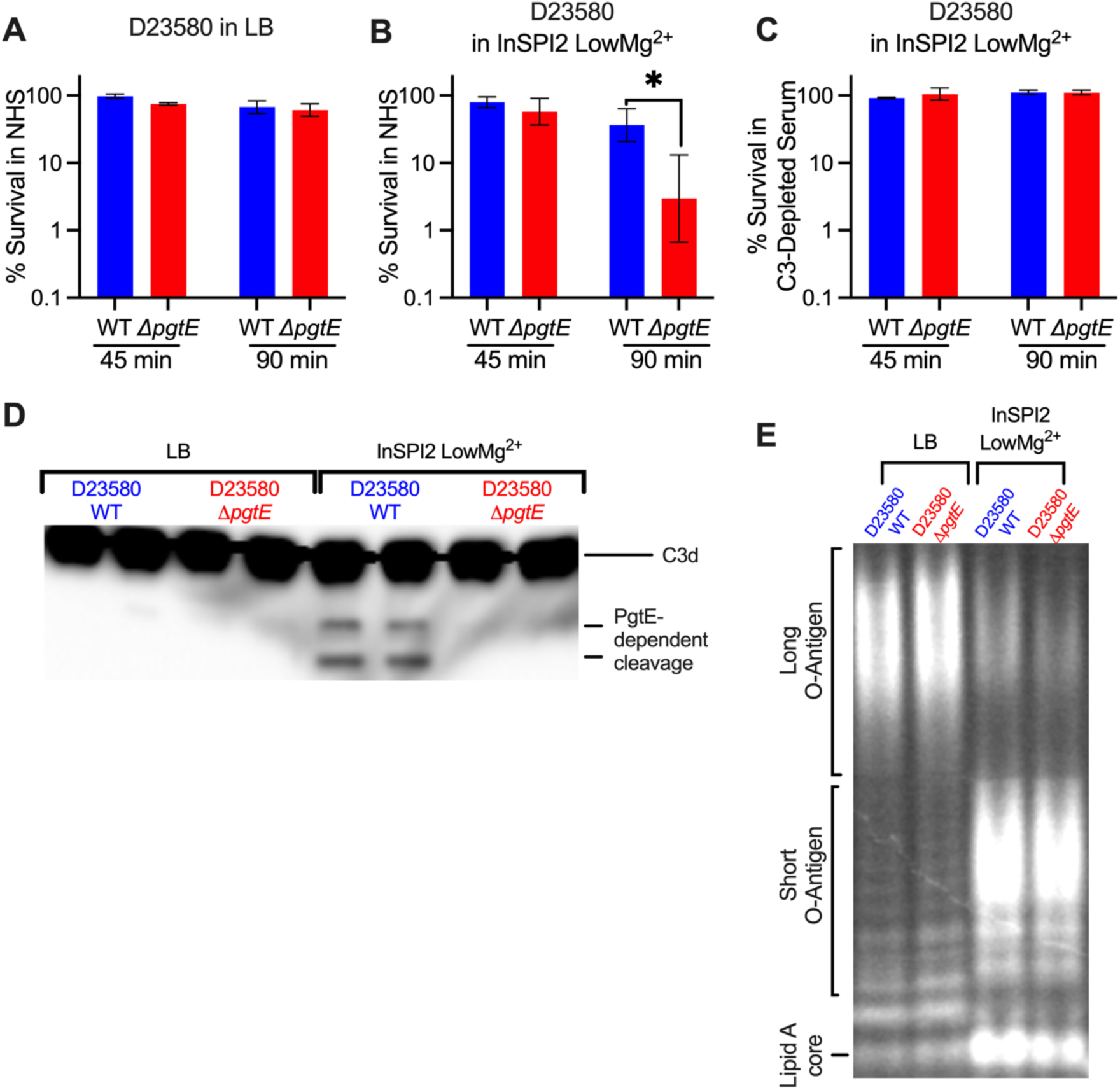
PgtE promotes survival of iNTS strain D23580 in serum when cultured in media mimicking the SCV luminal environment. (**A-C**) Serum killing assays were performed with smooth STm D23580 wild-type (WT) and an isogenic PgtE-deficient mutant (*ΔpgtE*). Strains were cultured overnight (**A**) in LB or (**B, C**) in InSPI2 LowMg^2+^ minimal media. STm at 10^6^ CFU/mL was then incubated with (**A, B**) 20% normal human serum (NHS) or (**C**) 20% C3-depleted human serum at 37 °C shaking at 300 rpm. CFUs were enumerated at 0 minutes, 45 minutes, and 90 minutes. % survival = (CFU at 45 minutes or 90 minutes / CFU at 0 minutes)*100%. (**A, C**) n = 2-3, (**B**) n = 6. Bar and error represent geometric mean and standard deviation. Data were analyzed by 2-way ANOVA followed by Sidak multiple comparison test. Adjusted p values from Sidak multiple comparison test: * p < 0.05. (**D, E**) D23580 WT and *ΔpgtE* were cultured overnight in (**Left**) LB or (**Right**) InSPI2 LowMg^2+^ minimal media. (**D**) After overnight culture, STm was lysed, supernatants were run on a 4-12% Tris-Glycine gel, and the gel was stained with Pro-Q Emerald 300 Lipopolysaccharide Gel Stain Kit to assess O-antigen chain length. (**E**) Alternatively, STm was then incubated with NHS for 8 hours. PgtE-dependent complement cleavage in supernatants was assessed by western blot analysis with anti-complement C3/C3b/iC3b/C3d antibody.

### PgtE expression enables STm to evade complement-mediated neutrophil killing

An important function of complement is to enhance neutrophil killing (48). To test whether PgtE-mediated complement cleavage enhances STm resistance to neutrophils, we cultured STm WT or the Δ*pgtE* mutant overnight in either LB (**Fig. 5A**) or InSPI2 LowMg^2+^ media (**Fig. 5B-C**) and infected neutrophils isolated from murine bone marrow. There was no difference in survival when the strains were grown in LB and either non-opsonized or opsonized with normal mouse serum (NMS) prior to the neutrophil infection (**Fig. 5A**). In contrast, when the strains were grown in InSPI2 LowMg^2+^ media and opsonized in NMS, STm WT survived significantly better than the Δ*pgtE* mutant in neutrophil killing assays (**Fig. 5B**). To assess if complement was the determinant factor in NMS for the difference in survival between STm WT and the Δ*pgtE* mutant, we opsonized the strains (cultured in InSPI2 LowMg^2+^ media) with serum from *C3^+/+^* or *C3^-/-^* littermate mice. Here, the survival defect of the Δ*pgtE* mutant in neutrophils was rescued to STm WT levels when the strains were opsonized in serum from *C3^-/-^* mice (**Fig. 5C**), indicating that PgtE enables STm to evade complement-mediated neutrophil killing.

**Figure 5.**
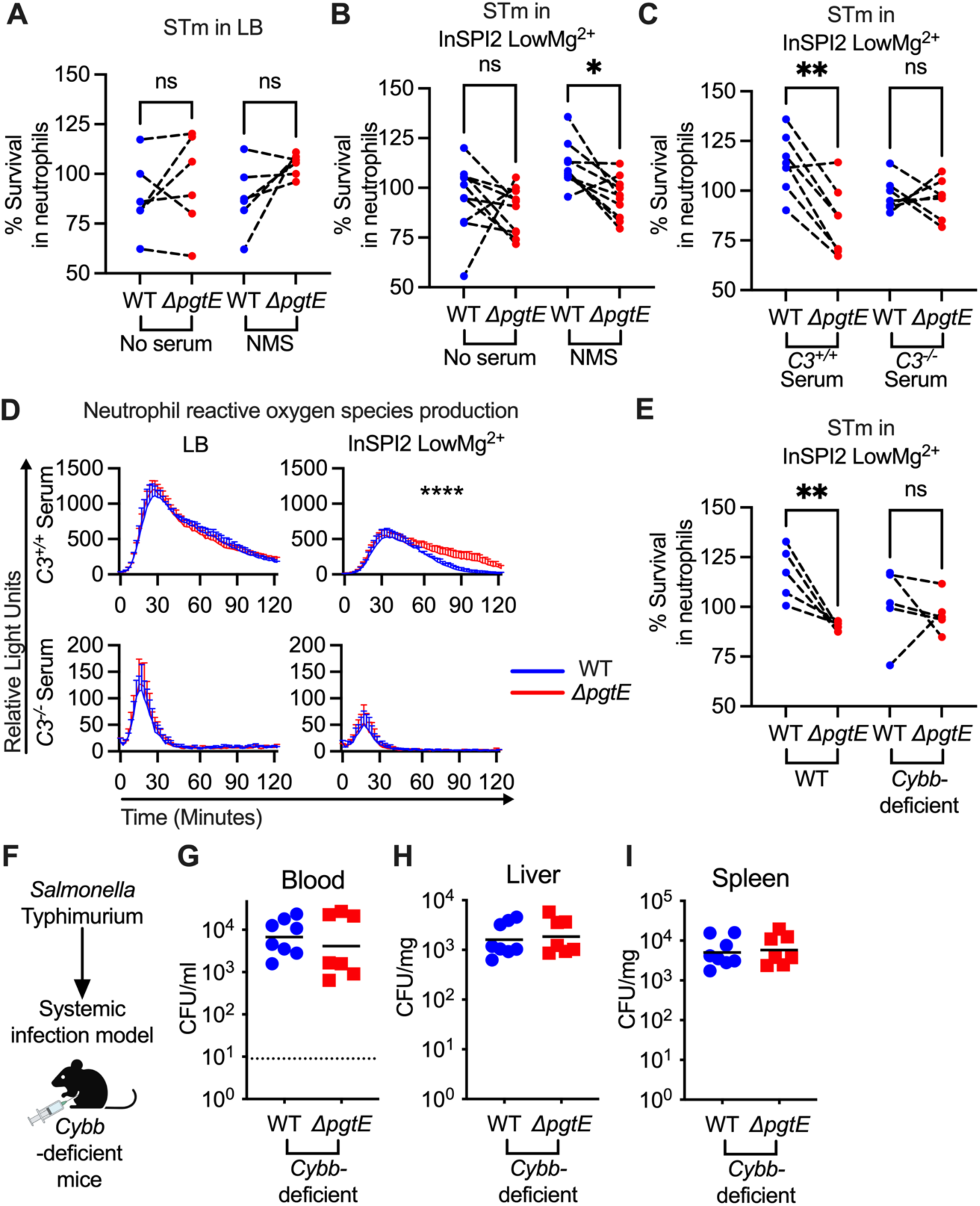
PgtE enhances STm survival in neutrophil killing assays and reduces complement-mediated neutrophil ROS response. Neutrophils were isolated (Stem Cell EasySep kit) from bone marrow of (**A-E**) C57BL/6 mice and (**E**) *Cybb*-deficient mice. For neutrophil killing assays, smooth STm IR715 wild-type (WT) and an isogenic PgtE-deficient (*ΔpgtE*) strain were cultured overnight in (**A**) LB or (**B-C, E**) InSPI2 LowMg^2+^ minimal media. STm was then (**A-B: Left**) not opsonized or (**A-B: Right, E**) opsonized with normal mouse serum (NMS). (**C**) Alternatively, STm was opsonized with serum from *C3^+/+^* and *C3^-/-^* littermates. (**A-C, E**) Neutrophils were then infected at an MOI = 10. STm CFU was enumerated 2.5 hours post-infection. % Survival in neutrophils = (CFU in wells with neutrophils at 2.5 hours/ CFU in control wells at 2.5 hours)*100%. (**D**) To determine neutrophil reactive oxygen species production, luminol assays were performed with STm cultured overnight in (**Left)** LB or (**Right**) InSPI2 LowMg^2+^ minimal media then opsonized with serum from (**Top**) *C3^+/+^* and (**Bottom**) *C3^-/-^* littermates. Neutrophils were infected at an MOI = 10. Relative Light Unit reads were performed every 2 minutes with a BioTek Synergy HTX. Error bars represent mean + SD from 3 biological replicates from 1 of 3 representative experiments. (**F-I**) 8-week-old *Cybb^X-/X-^* females or *Cybb^X-/Y^* hemizygous males were infected IP with 10^4^ CFU WT and *ΔpgtE* STm. Mice were euthanized 24 hours after infection and bacterial burden in the (**G**) blood, (**H**) liver, and (**I**) spleen was assessed. (**A-C, E**) N = 5-10 from 3-4 independent experiments. Symbols represent data with neutrophils from individual mice, bars represent the means. (**A-C, E**) Data were analyzed by One-way ANOVA Kruskal-Wallis test followed by Dunn’s comparison test. Adjusted p values from Dunn’s multiple comparison test: * p < 0.05, ** p < 0.01. (**D**) Data was analyzed by 2-way ANOVA. Time x Column Factor: **** p < 0.0001. (**D**) bar and error represent mean + SD. (**G-I**) Symbols represent data from individual mice, bars represent the geometric means. (**G**) Dotted line represents the limit of detection. (**G-I**) N = 7-8 from 2 independent experiments.

### PgtE disrupts C3-induced neutrophil ROS production, helping STm to evade ROS-dependent neutrophil killing

Complement enhances the neutrophil respiratory burst in response to STm (56, 57). To determine if PgtE disrupts C3-mediated reactive oxygen species (ROS) production by neutrophils, we performed a luminol assay with STm WT and the Δ*pgtE* mutant opsonized with serum from *C3^+/+^* or *C3^-/-^* mice (**Fig. 5D**). No differences were seen in neutrophil ROS production when the strains were grown in LB prior to opsonization with serum from *C3^+/+^*mice (**Fig. 5D**). In contrast, neutrophils infected with the Δ*pgtE* mutant exhibited prolonged ROS production compared to neutrophils infected with STm WT when the strains were cultured in InSPI2 LowMg^2+^ media and were opsonized with serum from *C3^+/+^*mice (**Fig. 5D**). Strains opsonized with complement-deficient serum induced lower levels of neutrophil ROS production, independent of PgtE expression (**Fig. 5D**). Thus, PgtE enables STm to evade the heightened ROS production that is triggered by C3 opsonization.

Next, we infected neutrophils isolated from wild-type or *Cybb-*deficient mice (**Fig. 5E**), which have defective ROS production (45). The Δ*pgtE* mutant exhibited comparable survival as STm WT in neutrophils from *Cybb*-deficient mice, indicating that PgtE promotes STm resistance to ROS-dependent neutrophil killing (**Fig. 5E**). When we infected *Cybb-*deficient mice intraperitoneally with STm WT or the Δ*pgtE* mutant (**Fig. 5F**), we recovered approximately 1-2 log more bacteria in comparison to WT mice in the blood, liver, and spleen (**Fig 5**; compare to Fig. 1). However, in *Cybb-*deficient mice, the Δ*pgtE* mutant was recovered to a similar level as STm WT in the blood (**Fig. 5G**), liver (**Fig. 5H**), and spleen (**Fig. 5I**). Thus, by disrupting C3-induced neutrophil ROS production, PgtE helps STm to evade ROS-dependent killing by neutrophils.

## DISCUSSION

Bacteremia is a major complication of NTS infection, and the mechanisms by which the pathogen evades host immune defenses are not fully understood. Here, we show that PgtE is a virulence factor that helps STm to overcome complement-mediated host defenses, survive in serum, and evade ROS-dependent neutrophil killing.

PgtE is an outer membrane protease that has been hypothesized to promote STm virulence through multiple mechanisms. For instance, PgtE expressed in rough strains of bacteria has previously been shown to promote adhesion to matrigel (35), suggesting a role for PgtE in enhancing invasion. PgtE also inactivates α2-antiplasmin while activating plasmin (40) and mammalian matrix metalloproteinase-9 (MMP-9) (37). Macrophages use plasmin and MMP-9 to migrate through tissues, and therefore PgtE was hypothesized to promote the dissemination of STm within infected macrophages (37, 40). Furthermore, STm can cleave cationic antimicrobial peptides (34), and multiple components of immune complement in a PgtE-dependent manner (36, 38, 39). Using immortalized human macrophage-like cells, a recent study showed increased localization of human bactericidal/permeability-increasing protein to SCVs containing PgtE-deficient STm, suggesting that PgtE promotes STm persistence in SCVs (47).

Collectively, studies with data generated mostly *in vitro* have proposed that PgtE enables STm to evade antimicrobial peptides and immune complement while promoting an intracellular lifestyle within macrophages. However, to our knowledge, no prior studies have linked these observations to *in vivo* phenotypes and specific components of host immunity, which requires the use of knock-out mice. Our results show that a STm Δ*pgtE* mutant is attenuated in the blood of wild-type mice, but fully rescued in *C3^-/-^* mice (**Fig. 1**), in mice treated with CVF (**Fig. 1**), and in *Cybb*-deficient mice (**Fig. 5**), thus demonstrating that PgtE promotes STm evasion of complement component C3 and ROS *in vivo*.

Identifying where and how PgtE plays a role *in vivo* was not trivial, as virulent STm has multiple virulence factors that modulate resistance to immune complement. For instance, long O-antigen chains confer serum resistance, but also sterically inhibit PgtE function (40, 58). Therefore, prior studies used rough STm and rough *E. coli* mutants when studying PgtE *in vitro* (36, 38, 39). Additional mechanisms of STm serum resistance include Rck and TraT, outer membrane proteins that confer serum resistance *in vitro* to either smooth or rough *E. coli* and *Salmonella* (31, 32, 59) by disrupting the complement membrane attack complex (MAC) (60). The many proposed functions of PgtE, by contrast, were observed in rough, avirulent strains.

Our study indicates that PgtE in fact does function *in vitro and in vivo* with fully virulent, smooth strains, albeit only after the physiologic O-antigen shortening that follows growth inside the SCV (40, 50, 51) (**Figs. 2, 4)**. A long O-antigen is a primary defense against an array of environmental insults, including immune complement activity. In environments where STm has a shortened O-antigen, such as in the SCV or having recently exited a phagocytic cell, PgtE likely represents a secondary line of defense to assist in protecting the more susceptible outer membrane.

Expression of *pgtE* and PgtE’s proteolytic function are enhanced in macrophages as well as in media that mimic the SCV lumen (40, 41, 49) (**Figs. 2, 4**). However, PgtE did not enhance STm survival in primary murine macrophages (**Fig. 2**), but did protect STm from C3-dependent serum killing (**Figs. 3, 4**). We obtained comparable results with the iNTS strain D23580 (clade ST313). When cultured in InSPI2 LowMg^2+^ media (mimicking the SCV lumen), strain D23580 exhibited reduced O-antigen length, cleaved C3 in a PgtE-dependent manner, and survived better in human serum (**Fig. 4**). These results are in agreement with a prior study that hypothesized that the increased expression of *pgtE*, due to a SNP in its promoter region, could enhance iNTS survival and dissemination (39).

A different study showed that, in response to serum exposure, multiple ST313 strains (including D23580), when cultured in LB, increased the expression of long O-antigen regulators but not of *pgtE*, *rck*, and *traT* (61). This suggests that when long O-antigen is present, STm continues to rely on the long O-antigen to resist complement killing. However, when the O-antigen is shortened (**Fig. 2, 4**), we demonstrate that PgtE defends against complement killing (**Fig. 3, 4**) and reduces neutrophil ROS production and killing (**Fig. 5**), thereby promoting bacteremia. Future studies will reveal whether PgtE also has other functions *in vivo*, and whether cleavage of other substrates contributes to STm pathogenesis.

## Abbreviations etc

BMDMs: bone marrow-derived macrophages
iNTS: invasive non-typhoidal *Salmonella*
NTS: Non-typhoidal *Salmonella*
STm: *Salmonella enterica* serovar Typhimurium
SCVs: *Salmonella*-containing vacuoles
SPI2: *Salmonella* Pathogenicity Island 2
InSPI2 LowMg^2+^: SPI2-inducing PCN media supplemented with low magnesium

## ACKNOWLEDGEMENTS

This work was funded by the NIH grant AI145325. Additional support was provided by AMED grant JP233fa627003, by the Chiba University-University of California-San Diego (UCSD) Center for Mucosal Immunology, Allergy, and Vaccines, and by the UCSD Department of Pediatrics. M.H.L. was supported by T32 DK007202 and F32 AI169989. JC was supported by NIH grant AI129992. LAK was supported, in part, by a Burroughs Wellcome PATH award. APL was supported by the NIAID Mucosal Immunology Studies Team (MIST). GTW was supported by NIH training grant T32AI007036. We would like to thank Ferric Fang for sending us the D23580 wild-type strain.

---

**Supplementary Table 1.**

| Designation | Genotype | Reference or Source |
| --- | --- | --- |
| ***Salmonella enterica* serovar Typhimurium** |  |  |
| IR715 | ATCC 14028s wild-type, spontaneous Nal<sup>R</sup> | Stojiljkovic et al, J. Bacteriol. 177(5):1357-1366 (1995) |
| JB10 | IR715 Δ*pgtE::tetRA* (Nal<sup>R</sup>, Tet<sup>R</sup>) | This study |
| ML27 | IR715 *pgtE*-FLAG (Nal<sup>R</sup>) | This study |
| LAKglmS | *SL1344 glmS::Ptrc-mCherryST::FCF* (Strep<sup>R</sup>, Cm<sup>R</sup>) | Knodler et al, Cell Host Microbe. 16(2):249-256 (2014) |
| LAKML2 | IR715 *glmS::Ptrc-mCherryST::FRT* (Nal<sup>R</sup>) | This study |
| D23580 | D23580 wild-type | Kingsley et al, Genome Research. 19:2279–2287 (2009) |
| SPN1113 | D23580 Δ*pgtE::tetRA* | This study |
| ***Escherichia coli*** |  |  |
| CC118 λ*pir* | F- *araD139* $\Delta$(*ara*, *leu*)7697 Δ*lacX74* *phoAD20 galE galK thi rpsE rpoB argE<sup>am</sup> recA1* λ*pir* | Herrero et al, J Bacteriol. 172(11):6557-67 (1990) |
| S17-1 λ*pir* | F- *recA thi pro rK- mK+ RP4:2-Tc::MuKm Tn7* λ*pir* | Herrero et al, J Bacteriol. 172(11):6557-67 (1990) |
| XL1-Blue | *recA1 endA1 gyrA96 thi-1 hsdR17 supE44 relA1 lac* [F *proAB lac<sup>q</sup>Z*ΔM15 Tn10 (Tet<sup>R</sup>)] | Agilent |
| DH5αMCR | F- *mcrA* $\Delta$(*mrr-hsdRMS-mcrBC*) ϕ80*lacZ*ΔM15 $\Delta$(*lacZYA-argF*)U169 *deoR recA1 endA1 phoA supE44*λ- *thi-1 gyrA96 relA1* | Gibco BRL |
| One Shot TOP10 | F- *mcrA* $\Delta$(*mrr-hsdRMS-mcrBC*) ϕ80*lacZ*ΔM15 Δ*lacX74 recA1 araD139* $\Delta$(*ara*, *leu*)7697 *galU galK rpsL*(Str<sup>R</sup>) *endA1 nupG* | Invitrogen |

**Supplementary Table 2.**

| Designation | Relevant characteristics | Reference or Source |
| --- | --- | --- |
| pCP20 | Ap<sup>R</sup>, temperature-sensitive, FLP recombinase system | Datsenko et al, Proc. Natl. Acad. Sci. USA. 97, 6640-6645 (2000) |
| pWSK29 | Ap<sup>R</sup>, MCS, *lacZa* | Wang et al, Gene. 100, 195-199 (1991) |
| pWSK29::*pgtE* | Ap<sup>R</sup> Tet<sup>R</sup>, pWSK29::*pgtE* (*pgtE* complementation) | This study |
| pWSK29::*pgtE*-D206A | Ap<sup>R</sup> Tet<sup>R</sup>, pWSK29::*pgtE*(D206A) (PgtE inactive allele) | This study |
| pRDH10 | Cm<sup>R</sup> Tet<sup>R</sup>, SacB (levansucrase: Sucrose sensitivity) | Kingsley et al, Applied and Environmental Microbiology, 1610-1618 (1999) |
| pRDH10::*pgtE*-FLAG | Cm<sup>R</sup> Tet<sup>S</sup>, SacB, pRDH10::*pgtE*-FLAG (*pgtE*-FLAG Tag) | This study |
| pCR-Blunt II-TOPO | Kan<sup>R</sup>, MCS | Invitrogen |
| pGP704 | Ap<sup>R</sup>, MCS, *oriR6K*, *mobRP4* | Miller et al, J. Bacteriology. 170(6):2575-2583 (1988) |
| pSPN23 | Ap<sup>R</sup> Tet<sup>R</sup>, pBluescriptII KS+::*tetRA* (*tetRA* cassette) | Raffatellu et al, Cell Host Microbe. 5(5):476-86 (2009) |
| pCRII::*pgtE*-LBRB | Kan<sup>R</sup>, pCR-Blunt II-TOPO::*pgtE*-LBRB (Δ*pgtE* cassette) | This study |
| pGP704::*pgtE*-LBRB | Ap<sup>R</sup>, pGP704::*pgtE*-LBRB (Δ*pgtE* cassette) | This study |
| pGP704::*pgtE*-LBRB::*tetRA* | Ap<sup>R</sup> Tet<sup>R</sup>, pGP704_*pgtE*_LBRB::*tetRA* (Δ*pgtE*::*tetRA* cassette) | This study |
| pP<sub>*pgtE*</sub>-gfp | Ap<sup>R</sup>, P<sub>*pgtE*</sub>-gfp<sup>mut3.1</sup> (*pgtE* transcriptional reporter plasmid) | This study |
| pMPM-A3ΔPlac | Ap<sup>R</sup>, P15A ori | Ibarra et al., Microbiology Apr;156(Pt 4):1120-1133 (2010) |

**Supplementary Table 3.**
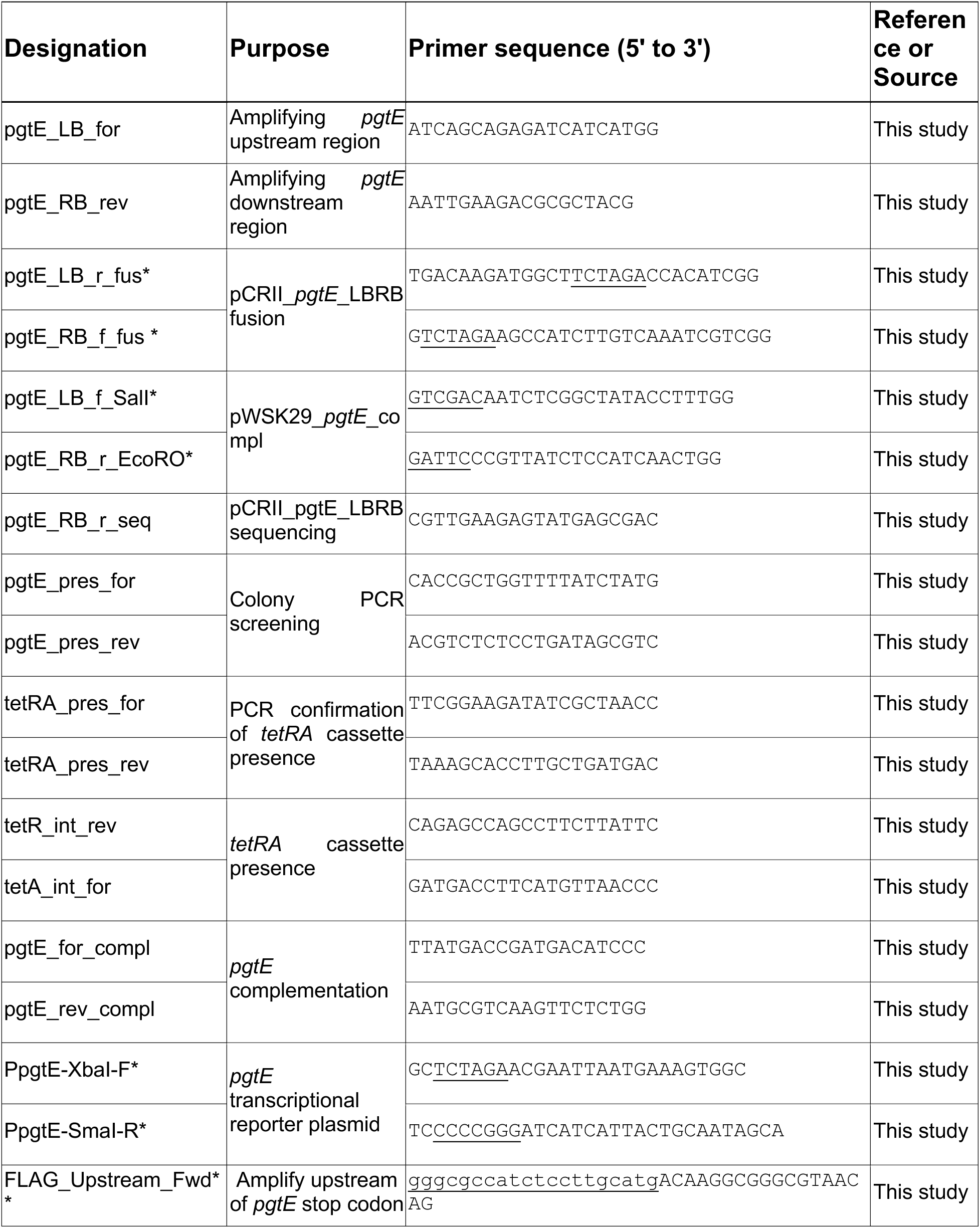
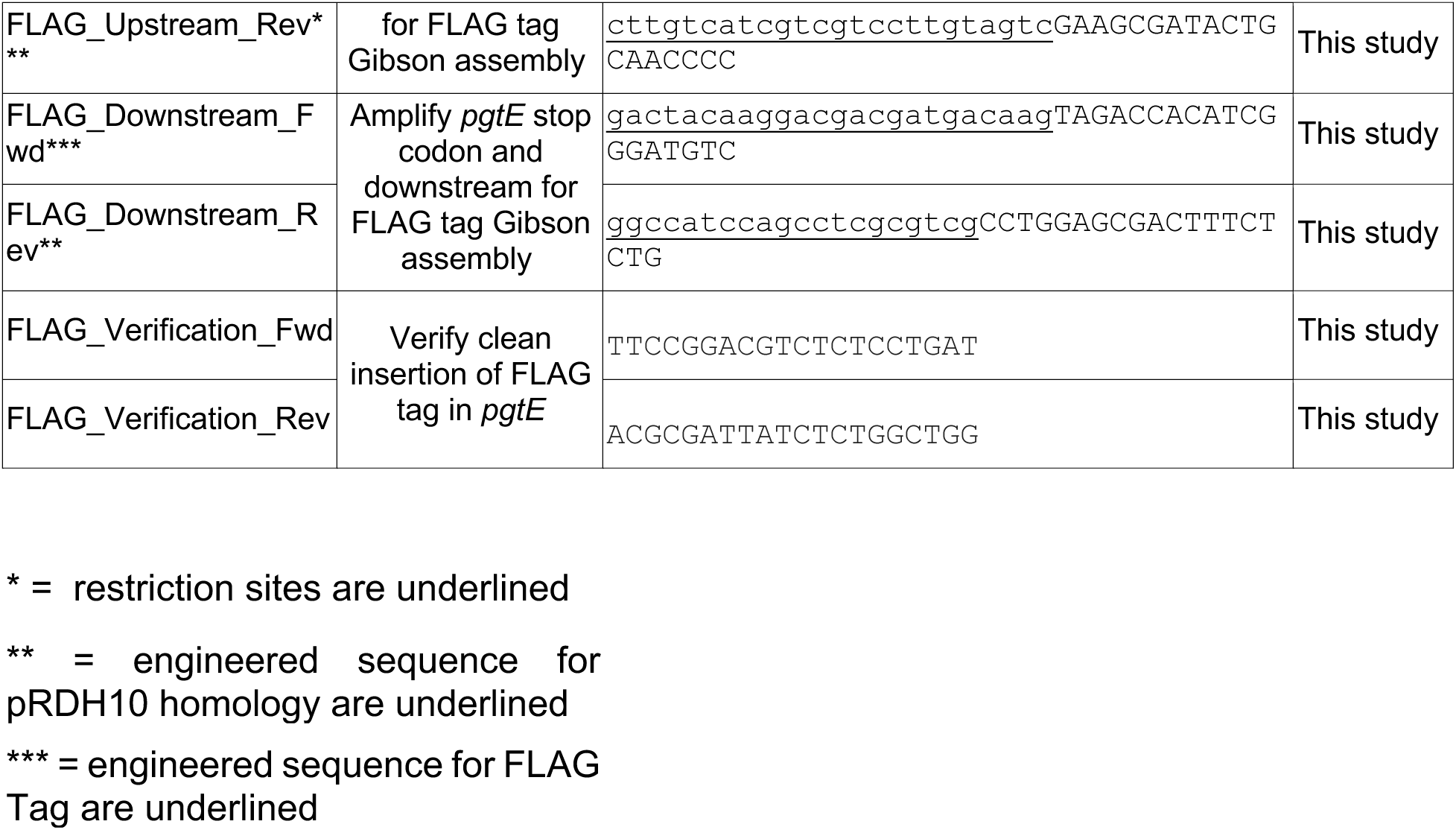

